# Exploring patterns of taxonomic, functional and phylogenetic β-diversity variation of Neotropical small mammals in a highly fragmented landscape

**DOI:** 10.1101/2022.08.09.503406

**Authors:** Wellington Hannibal, Nicolay Leme da Cunha

**Affiliations:** Laboratório de Ecologia e Biogeografia de Mamíferos, Universidade Estadual de Goiás, Quirinópolis, Goiás, Brazil; Grupo de Ecología de la Polinización, INIBIOMA, CONICET-Universidad Nacional del Comahue, San Carlos de Bariloche, Río Negro, Argentina

**Keywords:** abundance, geographical distance, NDVI, vegetation structure

## Abstract

Diversity can be partitioned in several components and dimensions that are affected in different ways by habitat loss and fragmentation. However, these partitions and dimensions are rarely investigated on human-modified landscapes. In this study, we investigated different partitions (Hill numbers) and dimensions (taxonomic [TβD], functional [FβD] and phylogenetic [PβD]) of small mammal β-diversity in a fragmented landscape of central Brazil using a multi-scale approach. TβD was estimated considering rare, common and abundant species. Tolerance to disturbed habitat, assessed via the traits “habitat use”, “tail length” and “use of vertical strata”, and trophic guild, defined by the “diet”, were used to estimate FβD. PβD was based on phylogenetic relatedness of the sampled species. The association between different partitions and dimensions of β-diversity with habitat and landscape attributes were investigated using Mantel tests. We found a significant positive effect of geographical distance on all partitions and dimensions of β-diversity. NDVI was the second most important variable affecting abundance based TβD, and all phylogenetic and functional β-diversity dimensions. Habitat characteristics, such as fallen logs and canopy cover were positively associated with all β-diversity dimensions. Our findings support the hypothesis that even in a highly modified landscape, small mammal’s β-diversity is determined by different environmental factors and spatial disposition of forest patches. However, the relatively higher importance of space appears to be related to dispersal limitation of this group.

## Introduction

Biological diversity is a topic of great interest for biologists and can have different connotations [1,2]. There are several ways to describe biological diversity, and among them the definition of three main components: alpha diversity (α) - diversity on a local scale, beta diversity (β) - the variation of species between locations, and gamma diversity (γ) - diversity on a regional scale [3] have been largely employed. Nevertheless, the traditional definition of β-diversity is dependent on α-diversity [3,4], leading to spurious results when researchers compare β-values of regions with different α-diversities [5]. Therefore, to avoid misinterpretation, α and β components should be ideally transformed into their number equivalents [6,7]; for example ^*q*^*D*_β_ for β-diversity, where *q*-number could determine a diversity measure’s sensitivity to rare or common species [5,8]. In this approach, species richness, Hill-Shannon diversity and Hill-Simpson diversity [6] are the three forms of Hill diversity most commonly used in ecological studies [8] and are generally known as the “Hill numbers”.

In a community dynamic viewpoint, β-diversity increases (heterogenization) when common species do not co-occur from some or all sites, or when new species arrive at some sites; and β-diversity decrease (homogenization) when rare, non-common species become extinct, or when formerly rare or absent species become widespread [9]. Variation on the trajectory of β-diversity can be caused by different effects of human disturbance [9–11]. Between these human disturbances, habitat loss has a consistently negative effect on biodiversity, while habitat fragmentation has been associated with the concept of habitat spatial heterogeneity, generally considered to have a positive influence on population and community-level ecological response [12,13]. In this scenario of habitat fragmentation, β-diversity should increase due mainly the effects of environmental variation among patches.

Nonetheless, the accrued evidence showed that the patterns of β-diversity in fragmented landscapes can result in homogenization or differentiation depending on the landscape heterogeneity and the spatial scale of analysis [14–17], highlighting the importance of isolation by distance on determining patterns of β-diversity variation [18]. Functional and phylogenetic β-diversity have been relatively less investigated on fragmented landscapes when compared to traditional taxonomic metrics [19], but despite less used, some studies show that a decrease in such dimensions of β-diversity was associated with land use intensification [20,21]. On the other hand, in a gradient of habitat complexity, the increase of all dimensions of β-diversity can be driven by β-replacement (TβD and FβD) or β-richness (PβD) [22]. So, ecological drives of biodiversity can differ among different biotas, scales and diversity facets [23]. Thus, decomposition of biodiversity into different dimensions may allow the identification of the main aspects of communities that are affected by forest conversion and fragmentation [24]. Therefore, in the abovementioned examples, habitat loss and fragmentation can lead to contrasting effects on β-diversity depending on the scale and dimension investigated.

Previous studies in the Neotropical region have shown that small mammals are good model organisms for testing the influence of landscape heterogeneity and habitat complexity on community ecology [25–30]. However, studies conducted in a multi-scale approach pointed contrasting scale-effects on small mammal β-diversity. For instance, in an Atlantic Forest fragmented landscape, small mammal β-diversity increased in small and isolated fragments [25], whereas, in Amazonian land-bridge islands, small mammal β-diversity was more strongly related to environmental variation (habitat quality) among sites than to spatial distance, patch scale and landscape scale [31]; further, in this ecoregion, all dimension of β-replacement decreased, while β-richness increased with forest area [32]. In the Brazilian Savanna, spatial configuration of the landscape and the extent, and quality of habitat strongly influence the rate of species turnover [30]. So, there is no consensus of the effects of landscape variation on structuring the Neotropical small mammals β-diversity, clearly demonstrating the need of further studies exploring different community dimensions to better understand the effects of habitat fragmentation on this important group of vertebrates.

In this study, we evaluated the response of species richness, Shannon’s entropy and Simpson’s dominance on TβD (multiplicative partitioning of Hill numbers) [5] to understand the relative importance of rare and dominant species on determining patterns of β-diversity in a highly fragmented landscape. We also investigated which are the main predictors of habitat (local) and landscape (regional) scales affecting the partitions of taxonomic, functional and phylogenetic β-diversity. Dispersal ability and habitat selectivity have been described as the main processes related to change on species composition [28], functional structure [29] and species turnover [30] of small mammal community in highly fragmented landscapes. Considering the relative low dispersal ability of small mammals [33–35], and the effect of habitat and landscape scale on structuring small mammal composition in Neotropical region [26,28,30,31], we would expect a positive effect of space, habitat and landscape quality on influencing all partitions and dimensions of β-diversity.

## Materials and Methods

### Study area

The study was conducted in southern Goiás State, central Brazil (18°25’ – 18°43’ S, 50°48’ – 50°22’ [28]), a highly fragmented landscape with about 13% of forest cover (https://mapbiomas.org/), dominated by semi-deciduous forest connected to riparian forest in a transitional region between Atlantic Forest and Cerrado ecoregion [36]. The climate is Tropical semi-humid - Aw (Köppen) with markedly dry season (April to September) and wet season (October to March), and mean annual temperature and mean annual rainfall around 23 °C and 1600-1900 mm, respectively [37].

### Sampling design

The sampling was carried out between January and December 2015. We captured small mammals, and quantified the vegetation structure and food resources in 24 trapping grids, distant 0.5 to 1 km within the same patch. Each trapping grid was composed of 20 trap stations, representing an area of 45 × 60 m (see, [28] for details about trapping grids). For landscape characterization, we estimated landscape metrics in 13 buffers of 1 km radius from the edge of sampled forest patches. The average forest patch size was 84.1 ha, with the smallest patch with 39.5 ha and the largest one with 142 ha. The average Euclidean distance among them was 22,588 m, being the closest distant 389 m and the farthest 51,978 m.

### Small mammal surveys

We captured small mammals using live traps (wire-cage traps and Sherman traps) and pitfall traps. We set Sherman and wire-cage traps in 16 trap-stations arranged in four linear transects, with 15-m intervals between the nearest trap stations. Each trap-station was composed by one wire-cage and one Sherman trap disposed on ground and understory (1.5 to 2 m height), alternatively. Pitfall traps was arranged in a perpendicular transect distant 15 m of each live trapping-grid, composed by four buckets (30 liters) connected by a fence of 0.8 m height (see, [28] for methodological detail). We made the captures under the collection license SISBIO nº 46985-1 and in accordance with guidelines provided by the American Society of Mammalogists [38].

### Habitat and landscape characterization

We used 23 variables to describe habitat and landscape characterization (S1 Data). For habitat characterization, we measured six variables related to vegetation structure (numbers of trees, shrubs, lianas, fallen logs, canopy cover and litter cover) and two variables that depict food resource availability (arthropods and fruits and seeds composition) in 10 selected trap stations of each trapping grid. We reduced the dimensionality of food resource availability using a Principal Coordinates Analysis (PCoA) separately for the matrix of arthropods and fruits-seeds resources. We associated the both matrices with the “horn” dissimilarity index, the one that presented higher variance recovery in relation to the original distances. As for the patch scale, we used four variables (i.e., forest area, core area, perimeter and NDVI), whereas for landscape scale, we used nine variables (i.e., mean isolation, water distance, shannon diversity, shape, mean perimeter area, core area, index core area, total edge and connectivity). The attributes related to patch and landscape scales were obtained via satellite images (methodological details can be found in [28]).

### Functional traits and phylogeny building

We selected four functional traits associated with tolerance to habitat disturbance: 1) habitat use [nominal trait: generalist or forest specialist], 2) tail length [quantitative trait in mm: arboreal species tend to be long tailed], 3) understory use [quantitative trait in %: based on the percentage of capture in the understory]; and trophic guild: 4) diet [multi-choice: insectivore, frugivore, granivore and omnivore, according to Annotated Checklist of Brazilian Mammals [41] (see, [29] for functional traits interpretation details). Traits were measured in the field from captured specimens or obtained from the literature [26,39–42].

We derived a phylogeny for our data set using the VertLife.org phylogeny subset on-line tool. We used as backbone the “Mammal’s birth-death node-dated completed trees (all 5911 species, set of 10k trees)” tree. Despite the criticisms regarding the use of synthesis-based phylogenies in evolutionary community studies, mostly because the relative low resolution and higher number of polytomies [43,44] demonstrated that this method of generating phylogenetic trees is sufficiently robust for community phylogenetic analysis.

### Data analysis

We initially tested for multicollinearity among all the 23 predictor variables described above using the Variance Inflation Factor (VIF) approach. For this, we used a stepwise procedure, where all data set was tested for collinearity, and when a variable showed VIF value above the threshold of 10, it was excluded and the procedure repeated with remaining variables until no variable was excluded. After such procedure, a total of 15 variables remained to be used as predictors in our models (S1 Table).

We used various distance-based metrics at the taxonomic, functional and phylogenetic dimensions to test for variation of β-diversity along the predictor variables selected above. For the taxonomic dimension, we partition the diversity of the metacommunity into β-components weighting different orders of diversity: species richness [q = 0], Shannon’s diversity [q = 1] and Simpson’s dominance [q = 2] (Hill numbers, [5-7]). For each Hill number, we did such partition for each pair of grids; thus, we estimate the β-diversity variation among all possible pairs of grids to generate a distance-based triangular matrix of β-diversity.

As for the functional dimension, we associated each pair of species by the eight continuous traits using the “Gower” distance. We then used this distance-based triangular matrix to estimate the Rao’s quadratic entropy [45], a measure of diversity in ecological communities accounting for species differences (functional or phylogenetic). We estimate two metrics of FβD: Dkl, which is the pairwise functional distance, and H, which is the Dkl standardized to account for within-community diversity. Both metrics return distance-based triangular matrix of FβD among pair of grids. For the phylogenetic dimension, we used the phylogenetic relationship of the captured species of small mammals to calculated the mean phylogenetic distance among all pairwise combinations of species co-occurring in a sample (MPD), which is a basal measure of the phylogenetic relatedness, and phylogenetic distance between each species and its nearest neighbor on the phylogenetic tree (MNTD), which is can be interpreted as a terminal metric of the phylogenetic relatedness of co-occurring species [46,47].

To test the association between each of the explanatory variables with the different dimensions of β-diversity, we used Mantel tests with 9999 permutations. For this, we used each of the β-diversity distance based on the triangular matrix as a response matrix, and a Euclidean distance-based on a matrix associating each sample unit by each predictor as a predictor matrix. We have also used a triangular matrix based on the Euclidean distance among grids to test for spatial autocorrelation. Moreover, given that our dataset is composed by 13 forest patches, being 12 of them with two grids, for testing the effect of the landscape metrics, we sampled one trapping grid per forest patch, thus reducing the number of sampling units to 13 to avoid pseudo-replication at a landscape scale. In the end, we did a total of 16 mantel tests for each β-diversity dimension described above. Finally, we tested for the spatial autocorrelation of each of the 15 predictor variables, and whenever we found an association between a β-diversity metric and any metric spatial autocorrelated, we did partial mantel tests in order to account for any potential confounding effect between explanatory variables.

We did all our analyses and graphics in R version 4.0.5 [48]. For calculating the VIF we used the package “*usdm*” [49]. To calculate the pairwise β-diversity at the taxonomic level, we used the function “*DivPart*” of the package “*entropart*” [50]. To associate our functional matrix based on “Gower” distance, we used the function “*vegdist*”, and to perform the Mantel tests, we used the “mantel” function, both from the package “*vegan*” [51]. To estimate the Rao’s quadratic entropy, we used the function “*raoD*”, and to calculate “MPD” and “MNTD” we used the functions “*mpd*” and “*mntd*”, respectively, all from the “picante” package [52].

## Results

### Small-mammal diversity

With an effort of 12,096 trap-night, we had 624 captures, resulting in a trap success of 5.2%. We captured 408 individuals (mean ± standard deviation (SD), 16.67 ± 8.76 individuals per trapping grid), belonging to 12 small mammal species (4.58 ± 1.50 species per trapping grid). The most common species were the arboreal marsupial *Gracilinanus agilis* (124 individuals, 20 trapping grids); the arboreal rodents *Rhipidomys macrurus* (55, 14), *Oecomys bicolor* (52, 16) and *Oecomys catherinae* (27, 6); the scansorial marsupials *Didelphis albiventris* (49, 20) and *Marmosa murina* (28, 8); and the terrestrial rodent *Calomys expulsus* (55, 13), comprising 96.1% of all individuals captured. The rarest species represented by less than 20 individuals were the terrestrial rodent *Calomys tener* (6 individuals, 4 trapping grid); the scansorial marsupial *Cryptonanus chacoensis* (5, 4); the scansorial rodent *Oligoryzomys mattogrossae* (2, 2); the terrestrial marsupial *Monodelphis kunsi* (2, 2); and the arboreal marsupial *Caluromys philander* (1, 1).

### Explanatory variables variation

For habitat characterization – vegetation structure and food resource availability, we found the following amplitude variation: “no. of shrubs” (min = 62, max = 1027), “no. of lianas” (min = 86, max = 418), “no. of fallen logs” (min = 3, max = 58), “canopy cover” (min = 41%, max = 98%) and “litter cover” (min = 85%, max = 99%). We considered the first two PCoA axes obtained from “arthropods” resource matrix (variance recovery, *r*^2^ = 0.85), and the first two PCoA axes from the “fruit-seed” resource matrix (*r*^2^ = 0.87) to represent “food resource availability” (S2 Table). Landscape characterization was based on patch and landscape metrics, according to following amplitude variation parameters: “forest area of focal patch” (min = 39.6, max = 142.3 ha), “perimeter length of focal patch” (min = 2749, max = 21136 m), “NDVI” (min = 0.2800, max = 0.3370), “patch isolation” (min = 99, max = 4102 m), “water distance” (min = 171.6, max = 1258.4 m) and “total edge of landscape” (min = 6410, max = 60766 m). Average values for all variables patch and landscape variables can be found elsewhere (S2 Table). Within the 15 predictor variables, we have only found evidence that “forest area” was spatially autocorrelated (Mantel test, r = 0.48, p < 0.001). All other variables showed no signal of spatial structure (S3 Table).

### Functional and phylogenetic variation

The small mammal community represented a range of different ecological traits, seven species were classified as habitat generalist (e.g., marsupials [*C. chacoensis, D. albiventris, G. agilis* and *M. kunsi*] and rodents [*C. expulsus, C. tener, O. mattogrossae*]), and five species were forest specialist (marsupials [*C. philander* and *M. murina*] and rodents [*O. bicolor, O. catherinae* and *R. macrurus*]). In relation to tail length, the shorter and longer tails were found in the terrestrial *M. kunsi* (36.1 mm) and scansorial *D. albiventris* (297.8 mm) marsupials, respectively. Thus, tail length of marsupials ranged from 36.1 to 297.8 (mean ± SD, 156 ± 87.8 mm), while the tail length of rodents ranged from 58.6 to 148.2 (96.8 ± 34.3 mm). The use of understory ranged from 0 (*M. kunsi, C. expulsus, C. tener* and *O. mattogrossae*) to 100% (*C. philander*), with little variation within each small mammal group (marsupials: min = 0, max = 100, 52.3 ± 41.7; rodents: min = 0, max = 93.1, 42.2 ± 47 frequency). We found no association between tail length and understory use (r = 0.44, *df =* 10, p = 0.15), showing that these traits are complementary to described use of vertical stratum. For trophic guild, the small mammals captured species were assigned to a diet category representing a combination of feeding guilds, such as: frugivore-omnivore (*C. philander* and *D. albiventris*) and insectivore-omnivore (*C. chacoensis, G. agilis, M. murina* and *M. kunsi*); while rodents were classified as: frugivore-granivore (*C. expulsus, C. tener* and *O. mattogrossae*) and frugivore-seed predator (*O. bicolor, O. catherinae* and *R. macrurus*). Categories were based on Paglia et al. [41].

The marsupial species captured in our study belong to the family Didelphidae, and are distributed into two subfamilies: Caluromyinae (i.e., *C. philander*) and Didelphinae that can be sub-divided in three tribes: Marmosini (*M. murina* and *M. kunsi*), Didelphini (*D. albiventris*) and Thylamyini (*C. chacoensis* and *G. agilis*). The six rodent species comprised the suborder Myomorpha, family Cricetidae, subfamily Sigmodontinae, belonging to three tribes: Oryzomyini (*O. bicolor, O. catherinae* and *O. mattogrossae*), Phyllotini (*C. expulsus* and *C. tener*) and Thomasomyini (*R. macrurus*). The phylogenetic and functional relationships of the captured small mammal community can be found in Fig 1.

**Fig 1:**
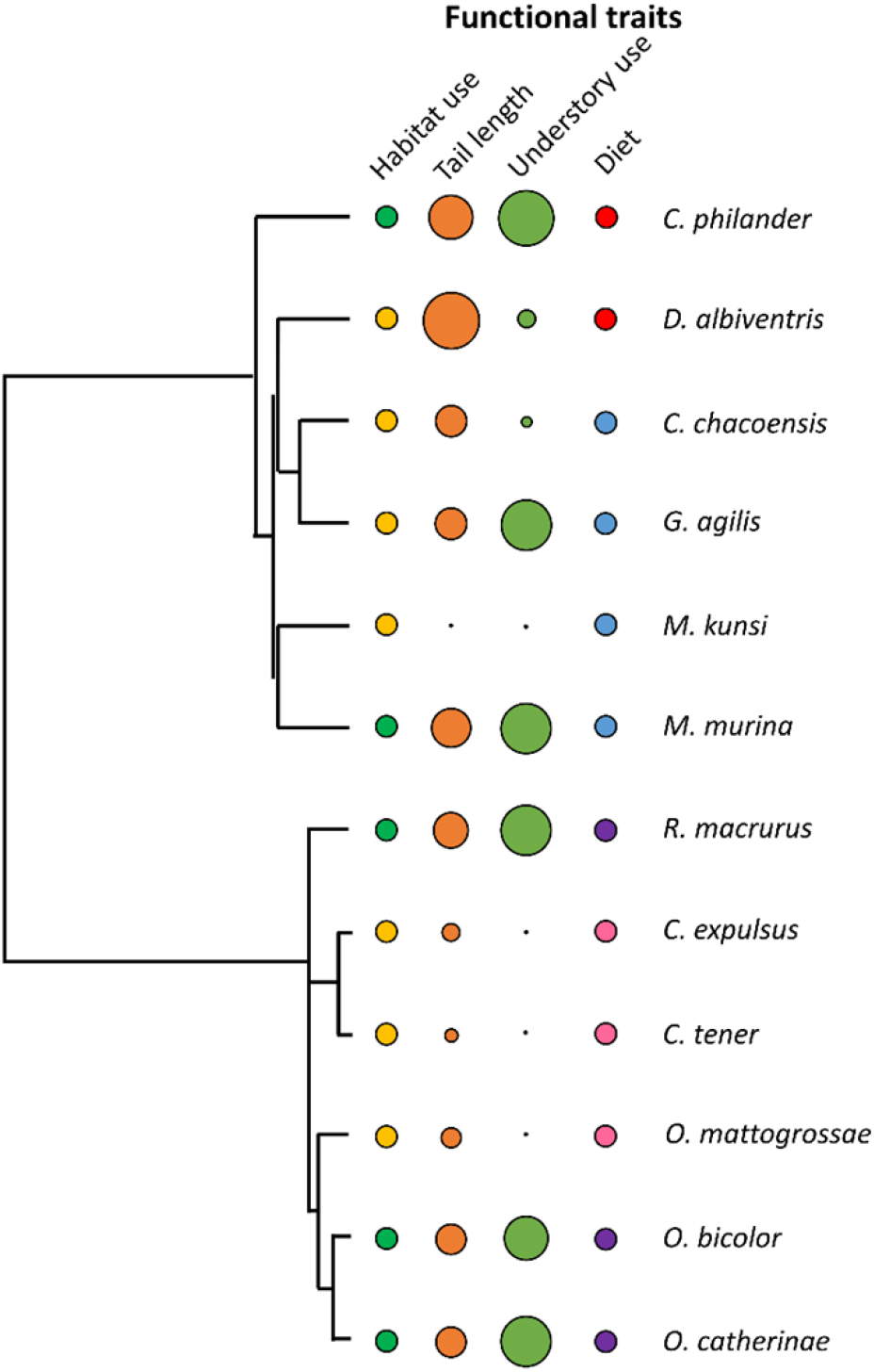
Phylogenetic hypothesis derived from the “VertLife.org phylogeny subset on-line tool”, using as backbone the “Mammal’s birth-death node-dated completed trees [all 5911 species, set of 10k trees]” and the functional traits used in our study: habitat use [nominal trait: generalist [orange circle] or forest specialist [green circle]], tail length [quantitative trait ranging from 31.6 to 297.8 mm], use of vertical strata [quantitative trait ranging from 0 to 100% and diet [multi-choice: frugivore-omnivore [red circle], insectivore-omnivore [blue circle], frugivore-granivore [pink circle] and frugivore-seed predator [purple circle]], based on relative capture success in traps set in the understory] [see, Hannibal et al. 2020 for functional traits interpretation details].

### Taxonomic β-diversity

The comparison of TβD variation with our predictor variables showed consistent results when weighting the species richness (q = 0) and Shannon entropy (q = 1). We found evidence of geographical distance (q = 0: r = 0.38, p = 0.008 [13 sample units]; q = 0: r = 0.32, p = 0.001 [24 sample units]; q = 1: r = 0.27, p = 0.047 [13 sample units]; q = 1: r = 0.34, p = 0.001 [24 sample units]; Fig 2a and b) on the patterns of β-diversity (Table 1). We have not found any other association with else predictor variables for these β-diversity metrics. For the β-diversity weighting the dominant species (q = 2), we found that geographical distance (r = 0.27, p = 0.042 [13 sample units]; r = 0.26, p = 0.004 [24 sample units]; Table 1, Fig 2c), “no. of fallen logs” (r = 0.22, p = 0.014; Table 1; Fig 2d) and “canopy cover” (r = 0.19, p = 0.038; Table 1, Fig 2e) were important predictors on determining the patterns of β-diversity turnover. Despite occurring all over the gradient, higher fallen logs frequency seems to determine the higher incidence of *C. tener, M. murina* and *R. macrurus*, and lower fallen logs frequency appears to associated to *G. agilis* and *C. philander* (S1 Fig). As for the canopy cover, most of the species were relatively abundant along the entire gradient, but some species like *M. kunsi, O. mattogrossae* and *C. philander* were exclusively found in grids with higher canopy cover, and *C. expulsus* was more frequently capture in low canopy cover sites (S2 Fig).

**Table 1.**
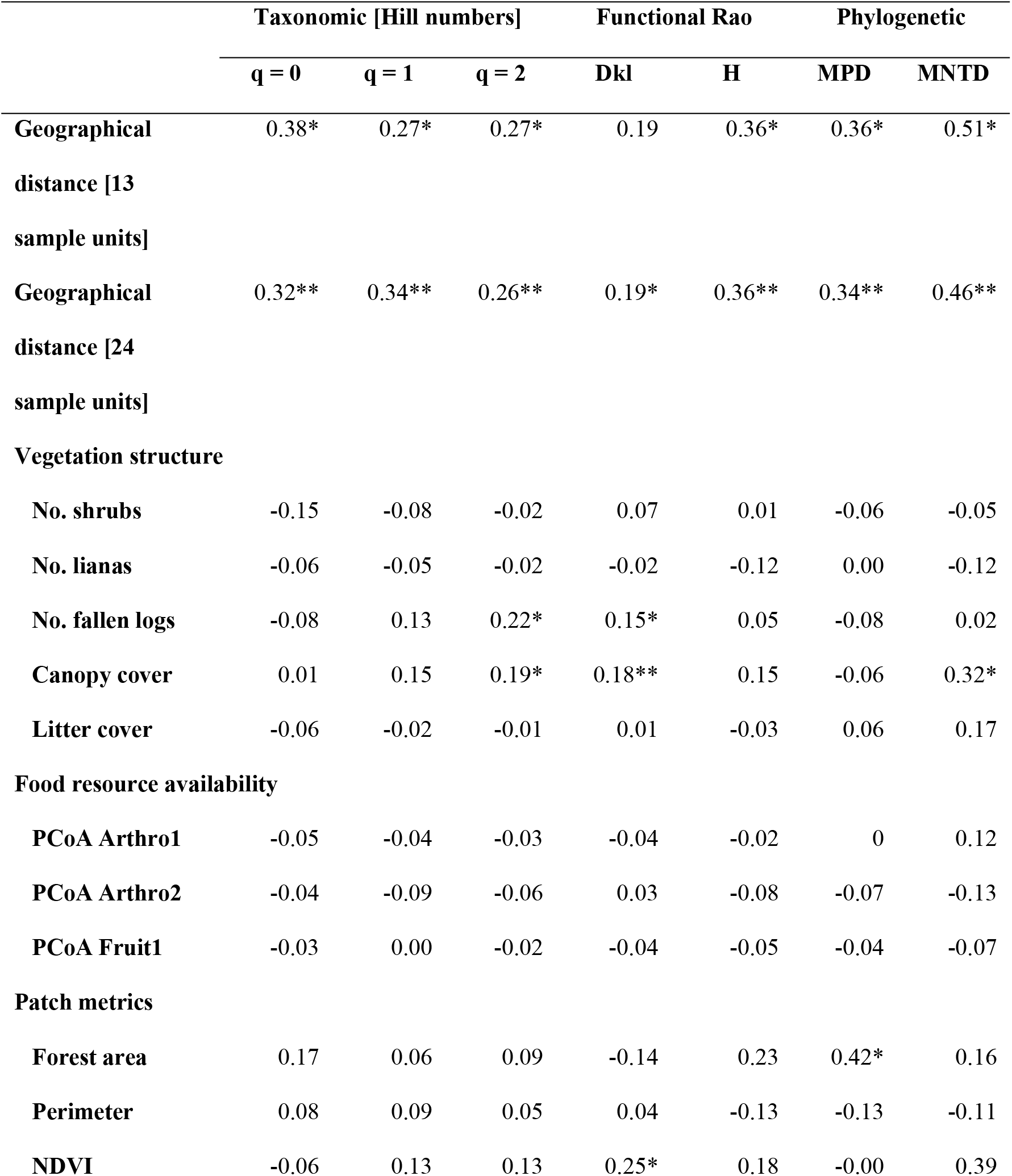
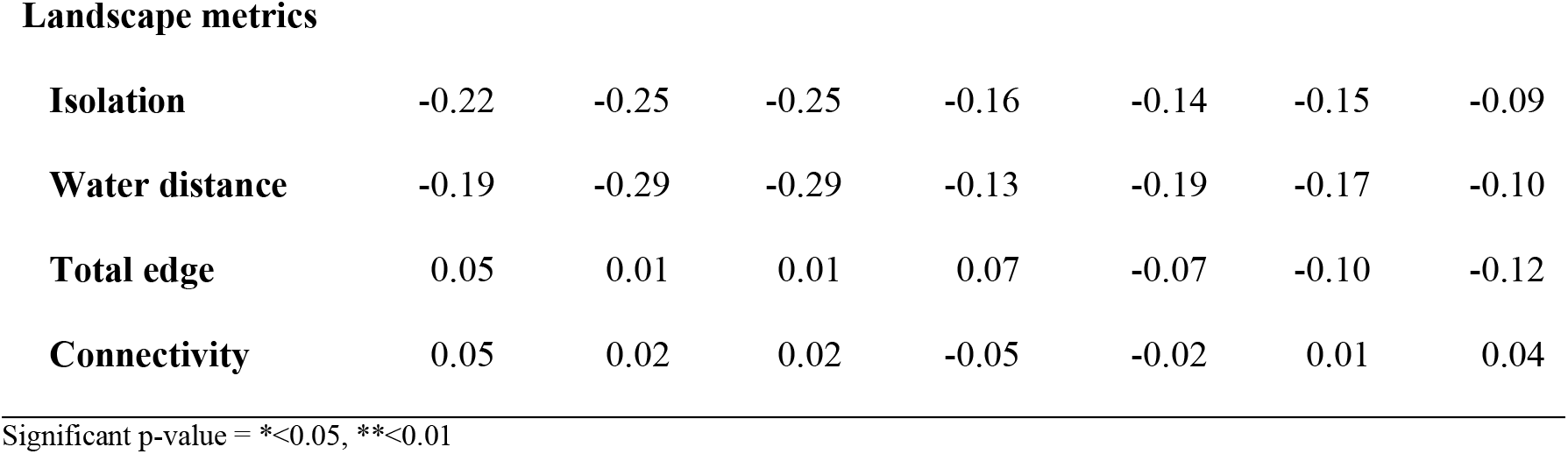
Pearson correlation between geographical distance, habitat characterization [vegetation structure and food resource availability] and landscape characterization [patch and landscape metrics] with taxonomic, functional and phylogenetic small mammal β-diversity. The significance of the relation was tested via Mantel-tests.

|  | Taxonomic [Hill numbers] |  |  | Functional Rao |  | Phylogenetic |  |
| --- | --- | --- | --- | --- | --- | --- | --- |
|  | q = 0 | q = 1 | q = 2 | Dkl | H | MPD | MNTD |
| **Geographical distance [13 sample units]** | 0.38* | 0.27* | 0.27* | 0.19 | 0.36* | 0.36* | 0.51* |
| **Geographical distance [24 sample units]** | 0.32** | 0.34** | 0.26** | 0.19* | 0.36** | 0.34** | 0.46** |
| **Vegetation structure** |  |  |  |  |  |  |  |
| **No. shrubs** | -0.15 | -0.08 | -0.02 | 0.07 | 0.01 | -0.06 | -0.05 |
| **No. lianas** | -0.06 | -0.05 | -0.02 | -0.02 | -0.12 | 0.00 | -0.12 |
| **No. fallen logs** | -0.08 | 0.13 | 0.22* | 0.15* | 0.05 | -0.08 | 0.02 |
| **Canopy cover** | 0.01 | 0.15 | 0.19* | 0.18** | 0.15 | -0.06 | 0.32* |
| **Litter cover** | -0.06 | -0.02 | -0.01 | 0.01 | -0.03 | 0.06 | 0.17 |
| **Food resource availability** |  |  |  |  |  |  |  |
| **PCoA Arthro1** | -0.05 | -0.04 | -0.03 | -0.04 | -0.02 | 0 | 0.12 |
| **PCoA Arthro2** | -0.04 | -0.09 | -0.06 | 0.03 | -0.08 | -0.07 | -0.13 |
| **PCoA Fruit1** | -0.03 | 0.00 | -0.02 | -0.04 | -0.05 | -0.04 | -0.07 |
| **Patch metrics** |  |  |  |  |  |  |  |
| **Forest area** | 0.17 | 0.06 | 0.09 | -0.14 | 0.23 | 0.42* | 0.16 |
| **Perimeter** | 0.08 | 0.09 | 0.05 | 0.04 | -0.13 | -0.13 | -0.11 |
| **NDVI** | -0.06 | 0.13 | 0.13 | 0.25* | 0.18 | -0.00 | 0.39 |
| **Landscape metrics** |  |  |  |  |  |  |  |
| **Isolation** | -0.22 | -0.25 | -0.25 | -0.16 | -0.14 | -0.15 | -0.09 |
| **Water distance** | -0.19 | -0.29 | -0.29 | -0.13 | -0.19 | -0.17 | -0.10 |
| **Total edge** | 0.05 | 0.01 | 0.01 | 0.07 | -0.07 | -0.10 | -0.12 |
| **Connectivity** | 0.05 | 0.02 | 0.02 | -0.05 | -0.02 | 0.01 | 0.04 |

677 Significant p-value = \* $<0.05$ , \*\* $<0.01$

**Fig 2.**
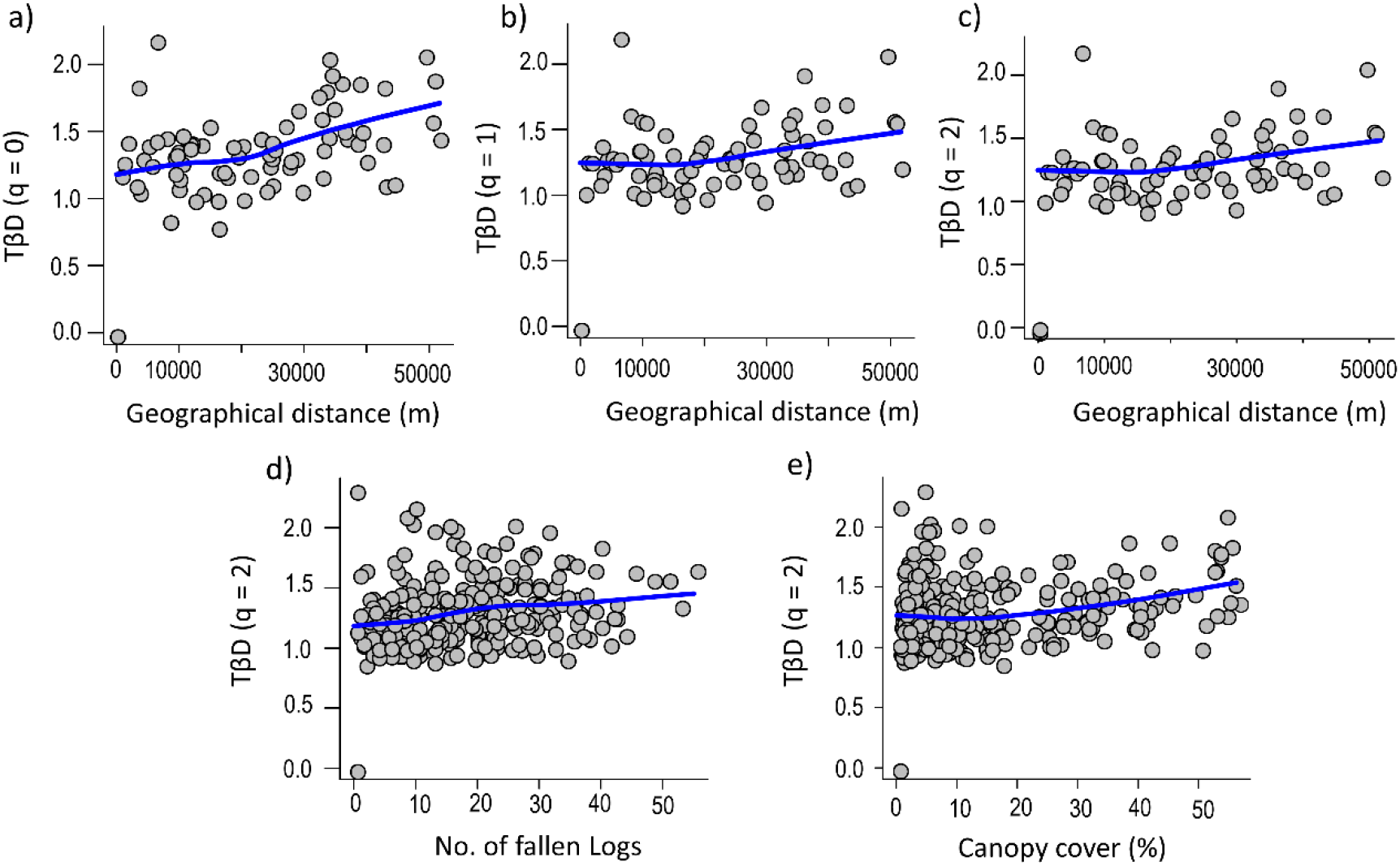
Correlation between small mammal taxonomic β-diversity and geographical distance [a-c], no. of fallen logs [d] and canopy cover [e] in a highly fragmented landscape in central Brazil. The Hill numbers associated represent rare [q = 0], common [q = 1] and abundant [q = 2] species, respectively. The tendency line is fitted using Locally Estimated Scatterplot Smoothing [LOESS].

### Functional and phylogenetic β-diversity

The functional turnover represented by Dkl was positively associated with “no. of fallen logs”, “canopy cover” and “NDVI” (r = 0.15, p = 0.030; r = 0.18, p = 0.004; r = 0.25, p = 0.018, respectively; Table 1, Fig 3a-c). In the other hand, the standardized H was related only to geographical distance (r = 0.36, p = 0.013; Table 1, Fig 3d). Higher values of Dkl and H were associated with species like *O. mattogrossae, D. albiventris, C. chacoensis* and *O. bicolor*; species with a wide range of variation in their traits related to use of habitat, tail length, and diet, which have determined such high values of functional diversity variation (S3 Fig).

**Fig 3.**
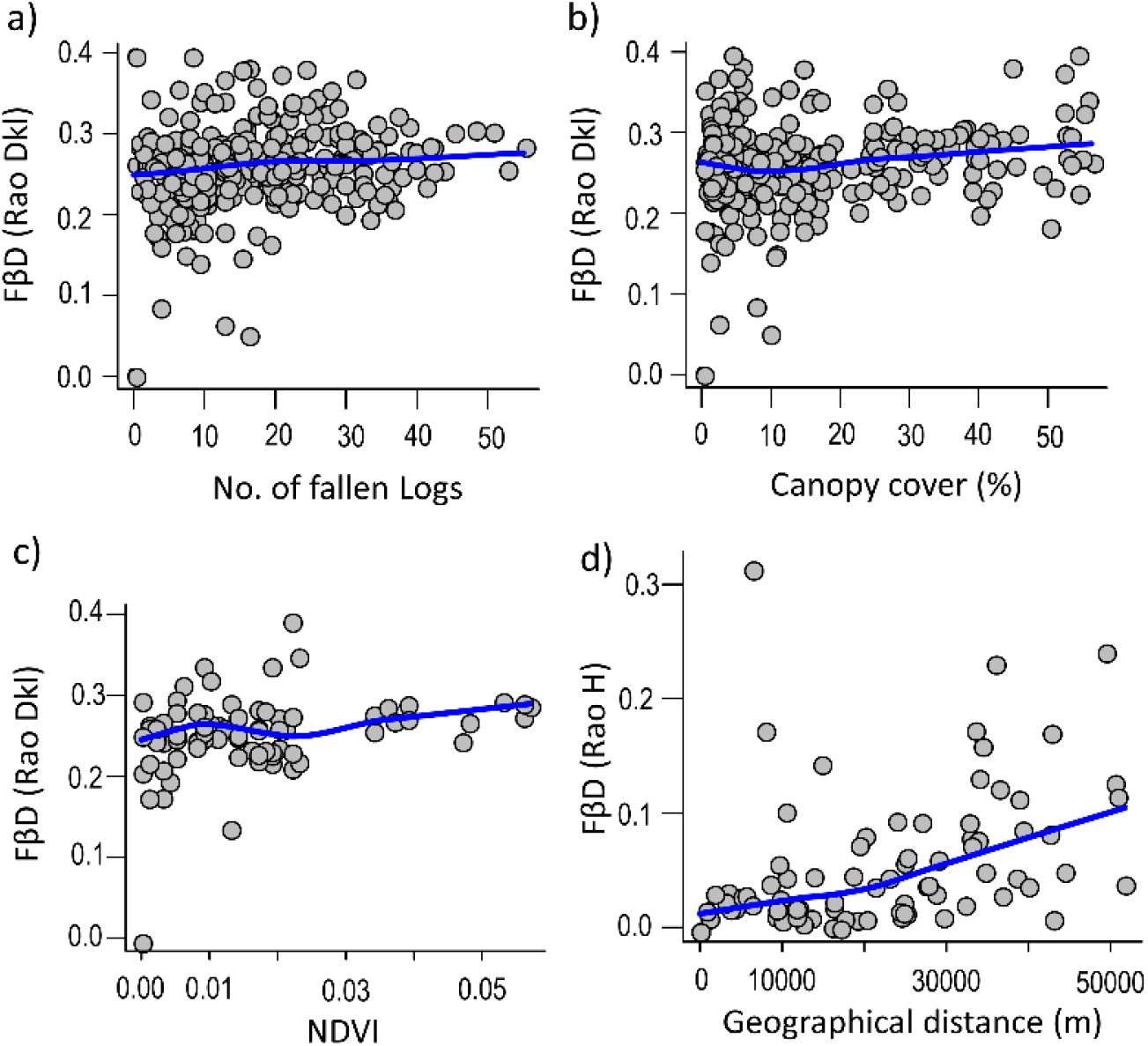
Correlation between small mammal functional β-diversity and no. of fallen logs [a], canopy cover [b], NDVI [c] and geographical distance [e] in a highly fragmented landscape in central Brazil. The tendency line is fitted using Locally Estimated Scatterplot Smoothing [LOESS].

The PβD represented by MPD showed a positive association with geographical distance and “forest area” (r = 0.36, p = 0.006 [13 sample units]; r = 0.34, p = 0.001 [24 sample units]; r = 0.42, p = 0.017; Table 1, Fig 4a and b). The effect of “forest area” is consistently significant even when accounting for the variability captured by the geographical disposition of the grids (Partial mantel, r = 0.31, p = 0.04). The MNTD showed significant associations with geographical distance and “canopy cover” (r = 0.51, p = 0.007 [13 sample units]; r = 0.46, p = 0.001 [24 sample units]; r = 0.32, p = 0.030; Table 1, Fig 4c and d). Marsupials, Marmosini tribe (*M. murina* and *M. kunsi*) and arboreal rodents, tribes Oryzomyini (*O. bicolor*) and Thomasomyini (*R. macrurus*) were more frequent in habitats with high “canopy cover” values (S2 Fig).

**Fig 4.**
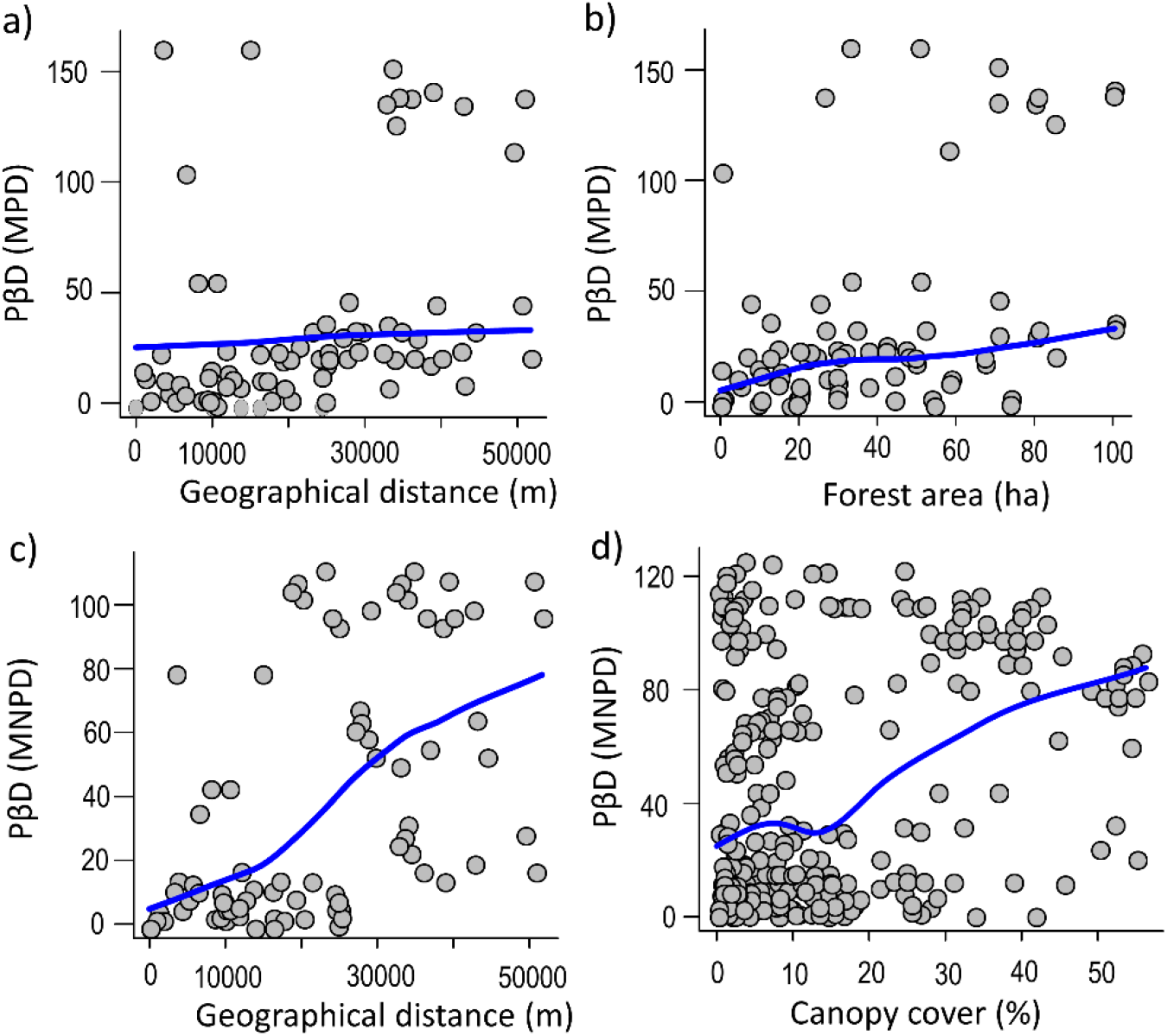
Correlation between small mammal phylogenetic β-diversity and geographical distance [a and c], forest area [b] and canopy cover [d] in a highly fragmented landscape in central Brazil. The tendency line is fitted using Locally Estimated Scatterplot Smoothing [LOESS].

## Discussion

### Overview

Our findings revealed that *i*) small mammal communities sampled in this highly fragmented landscape Neotropical region were dominated by commonly found marsupials and rodent’s species with arboreal-scansorial habits. *ii*) In spite of being highly fragmented, our landscape presented a wide variation of habitat quality, mostly driven by vegetation structure, food resource and landscape attributes, which harbor functionally and phylogenetically diverse small mammal communities in central Brazil. *iii*) Geographical distance was the main variable correlated to all partitions and all dimensions of β-diversity. Habitat quality gradient was an important predictor for TβD (partition of dominant species), and FβD and PβD. We discuss below such findings in the light of dispersal limitation and environmental filters on driving the patterns of β-diversity in small mammals of Central Brazil.

### Taxonomic β-diversity

Geographical distance was positively correlated with all partitions of TβD, corroborating our initial hypothesis. Marsupials and small rodents have a low dispersal ability [33,35,51,52] that limit the continuous replacement of small mammal species between distant localities. Further, small mammal communities in the Cerrado domain have been discussed elsewhere to be mainly driven by dispersal limitation and habitat selectivity [28,30]. Geographical distance also has been the most important predictor of small mammal dissimilarity in Atlantic Forest [52]. This finding can be explained by the idea that fragmented landscapes are hyper-dynamically influenced by environmental heterogeneity [12]. Small mammals are able to disperse between relatively closer fragments (∼ 485 m in average, according to [55]), but homing behavior can be much higher for larger species like *Philander frenatus*, which crossed an area of 1050 m of fragmented landscape in the Atlantic Forest [56]. In our study, although the landscape is mainly composed by homogeneous semideciduous forest fragments, strong heterogeneity at small scales can be found between forest patches (e.g., vegetation structure). Such variability may reflect different levels of habitat quality, which certainly influence the potential source pools and species establishment for each patch with different degrees of isolation.

When weighting abundant species (q = 2 Hill number), TβD was also positively associated to habitat quality (e.g., fallen logs and canopy cover). We captured an expressive number of individuals in such highly fragmented landscape of central Brazil, within these only *C. philander, M. kunsi* and *O. mattogrossae* were rarely recorded in our study (1 to 2 individuals). On the other hand, 96% of the captured individuals - seven of the twelve captured species - occurred in 6 to 20 trapping grids. This finding may be caused by the variation in vegetation structure between different sampling sites, which by consequence determine abundance patterns for the most common species. Thus, the higher values of TβD based on the abundance of species of small mammals can be resulted from the higher variability in small mammal community structure along this habitat quality gradient. In the Cerrado domain, the abundance (not richness) of small mammals is known to be correlated with different types of vegetation cover (e.g., herbaceous, shrub, and tree density) [57], a variable that is commonly associated with habitat heterogeneity and complexity [58].

### Functional and phylogenetic β-Diversity

In overall terms, the small mammals communities sampled in our study is relatively species poor when compared to other communities of the same group in Central Brazil [59], a pattern that seems to related to the elevated habitat loss in the region. Large bodied rodents that are specialists in forested habitats (e.g., tribe Oryzomyini [*Hylaemys megacephalus*], Family Echimyidae, Eumysopinae [*Proechimys longicaudatus* and *P. roberti*]) generally represent the first functional group and respective phylogenetic lineages to disappear in highly fragmented landscapes [25,60]. Such group is intimately associated with continuous forests and landscapes with high percentual of vegetation cover [25,27,39,60,61]. In the other hand, species of marsupials (e.g., *G. agilis* e *C. chacoensis*) and rodents (e.g., *C. expulsus* e *C. tener*) that have a broad ecological niche and are able to use both forested and open field environments, as present an opportunistic feeding habitat [29,60], commonly represent the species that are capable to be successful even in such disturbed environments. Small sized rodent species that use the understory (e.g., *O. bicolor* e *R. macrurus*) seems also to be weakly affected by habitat fragmentation in the region, perhaps the smaller scale of niche requirements of these species protected them of the negative effects of habitat fragmentation [29]. However, despite high co-occurrence of such resistant species between forest fragments in the region, variation in abundance patterns are common for these species, which may be related to different niche requirements and responses to habitat modification [29].

Despite the relatively modest phylogenetic and functional variation of our sampled small mammals’ communities, we found a turnover for these community dimensions in relation to geographical distance and habitat quality gradient (i.e., no. fallen logs and canopy cover). In the Amazonian habitat, FβD and PβD of bat communities were highest between continuous forest and *campinarana*, two highly contrasting environments, and these patterns in β-diversity were associated with functional richness and lineage richness differences, respectively [22]. An study with bird and ant assemblages, in Atlantic Forest and Pampas Grassland showed that land uses and biomes seems to promote assemblage differentiation in traits and lineages that occurred in anthropogenic habitats, further in this landscape both animal groups were similarly sensitive to changing in vegetation structure [63]. Therefore, habitat quality seems to be an important driver of functional and phylogenetic turnover in biological communities in the Neotropics independent of the biological group or sampled region. Nevertheless, understanding what are the most important traits and functional lineages that readily respond to habitat disturbance, as what are the most resilient ones that persist in disturbed fragments, are of utmost importance for providing information for conservation initiatives.

Little is known about the dispersibility of small mammals among forest patches in the Neotropics [55]. Despite reasonable to expect, a positive relation between body size and dispersiveness is not always the rule for this group of mammals. For instance, the mid-bodied size nocturnal marsupial *P. frenatus* (400-600 g) has a relatively small living area (2.8 ha). Thus, even being a larger species for a small-mammal, the apparent inability of this species to occupy different types of habitats (e.g. less forested patches or open fields), may limit its dispersion to equivalent nearby fragments, letting this species susceptible to local extinctions [64], which may similarly be the case for other large bodied terrestrial small rodents that was expected to be reported in our study, but were not captured [e.g. *Cerradomys scotti, C. maracajuensis, C. marinhus, C. subflavus, P. longicaudatus, P. roberti*]. The capacity of using the landscape matrix as a secondary habitat, confers more flexibility to overcome the negative effects of habitat fragmentation [65,66]. Therefore, the prevalence of mostly generalist’s species that can occupy different portions of the landscape found here may explain the low FβD e PβD in the sampling region.

## Conclusion

To the best of our knowledge, this is the first study that shows the effect of geographical distance, habitat and landscape variation on all partitions and dimensions of small mammals’ β-diversity. Considering that we have a depleted small mammal community due to high habitat loss in central Brazil, we conclude that only abundance-weighted β-diversity values responded to the predictors of habitat quality. In this sense, a homogeneous environment results in poor communities, where abundance was more important than richness, therefore, studies that seek to investigate the effect of habitat loss and fragmentation on biodiversity need to consider species abundance and not just species richness. Abundance was also a more important attribute than richness in anuran assemblages in Pantanal, a region naturally disturbed by floods in South America, since most species co-occurred in all sampled ponds but varied in their abundance [67]. Thus, we summarize that taxonomic, functional and phylogenetic replacement of the small mammal communities in the fragmented landscape of central Brazil is determined by dispersal limitation and habitat selection, driven by species-specific responses in the communities’ arrangements.

## Acknowledgments

We thank to the colleague of the Laboratory of Ecology and Biogeography of Mammals for field assistance.

## Supporting Information

**S1 Table.** VIFs of the remained variables after stepwise procedure, based on 23 variables that characterized habitat and landscape scales.

| Variables | VIF |
| --- | --- |
| No. shrubs | 3.196 |
| No. lianas | 2.673 |
| No. fallen logs | 3.750 |
| Canopy cover | 2.594 |
| Litter cover | 2.741 |
| PCoA Arthro1: Arthropods resources according to axis 1 of PCoA | 2.464 |
| PCoA Arthro2: Arthropods resources according to axis 2 of PCoA | 3.561 |
| PCoA Fruit-Seed1: Fruit-seed resources according to axis 1 of PCoA | 2.620 |
| Forest area of patch [ha] | 3.184 |
| Perimeter of focal patch [m] | 8.916 |
| NDVI | 3.557 |
| Isolation: mean isolation of the five nearest patches [m] | 1.957 |
| Water distance: distance to water course [m] | 3.780 |
| Total edge in the landscape [m] | 5.172 |
| Connectivity [connected *versus* isolated patch, considering riparian forest] | 4.424 |

**S2 Table.** Habitat, patch and landscape metrics that characterization fragmented landscape studied.

| Parameter | Type of parameter | Description | Range | Mean ± SD |
| --- | --- | --- | --- | --- |
| No. shrubs | Vegetation structure | Number of shrubs in trapping-grid | 62-1027 | 228.60±193.01 |
| No. lianas | Vegetation structure | Number of lianas in trapping-grid | 86-418 | 224.27±80.04 |
| No. fallen logs | Vegetation structure | Number of fallen logs in trapping-grid | 3-58 | 33.63±14.41 |
| Canopy cover | Vegetation structure | Canopy cover in trapping-grid | 41-98% | 85.92±14.70 |
| Litter cover | Vegetation structure | Litter cover in trapping-grid | 85-99% | 95.55±3.32 |
| PCoA Arthro1 | Food resource | PCoA axis 1 of arthropods in trapping-grid | - | - |
| PCoA Arthro2 | Food resource | PCoA axis 2 of arthropods in trapping-grid | - | - |
| PCoA Fruit-Seed1 | Food resource | PCoA axis 1 of fruit and seed in trapping-grid | - | - |
| PCoA Fruit-Seed2 | Food resource | PCoA axis 2 of fruit and seed in trapping-grid | - | - |
| Forest area | Patch metric | Total area of sampled patch [ha] | 39.6-142.3 | 81.52±36.23 |
| Perimeter | Patch metric | Perimeter of sampled patch [m] | 2749-21136 | 6189.2±4733.7 |
| NDVI | Patch metric | Normalized difference vegetation index | 0.28-0.33 | 0.32 ±0.01 |
| Isolation | Landscape<br>metric | Mean distance of the 4<br>nearest patches [m] | 99-4102 | 1603.1±1311.5 |
| Water<br>distance | Landscape<br>metric | Distance of trapping grid to<br>water course [m] | 171.6-1258.4 | 628.7±344.7 |
| Total edge | Landscape<br>metric | Total edge measured in the<br>landscape [m] | 6410-60766 | 23679.9±15560.5 |

**S3 Table.** Correlation between predictors variables and spatial distribution of the sampled forest patches.

| Group | Method | Statistic | N | P.value |
| --- | --- | --- | --- | --- |
| No. shrubs | Pearson | 0.0666 | 276 | 0.315 |
| No. lianas | Pearson | -0.0088 | 276 | 0.509 |
| No. fallen logs | Pearson | 0.0312 | 276 | 0.317 |
| Canopy cover | Pearson | 0.0759 | 276 | 0.205 |
| Litter cover | Pearson | 0.0167 | 276 | 0.401 |
| PCoA Arthro1 | Pearson | 0.0229 | 276 | 0.292 |
| PCoA Arthro2 | Pearson | -0.1020 | 276 | 0.837 |
| PCoA Fruit-Seed1 | Pearson | -0.0567 | 276 | 0.752 |
| Forest area | Pearson | 0.4495 | 78 | 0.007* |
| Perimeter | Pearson | -0.1249 | 78 | 0.745 |
| NDVI | Pearson | -0.0000 | 78 | 0.458 |
| Isolation | Pearson | -0.0210 | 78 | 0.502 |
| Water distance | Pearson | -0.0370 | 78 | 0.572 |
| Total edge | Pearson | -0.0896 | 78 | 0.721 |
| Connectivity | Pearson | -0.0836 | 78 | 0.831 |

### Supporting Information

#### Figures

**S1 Fig.**
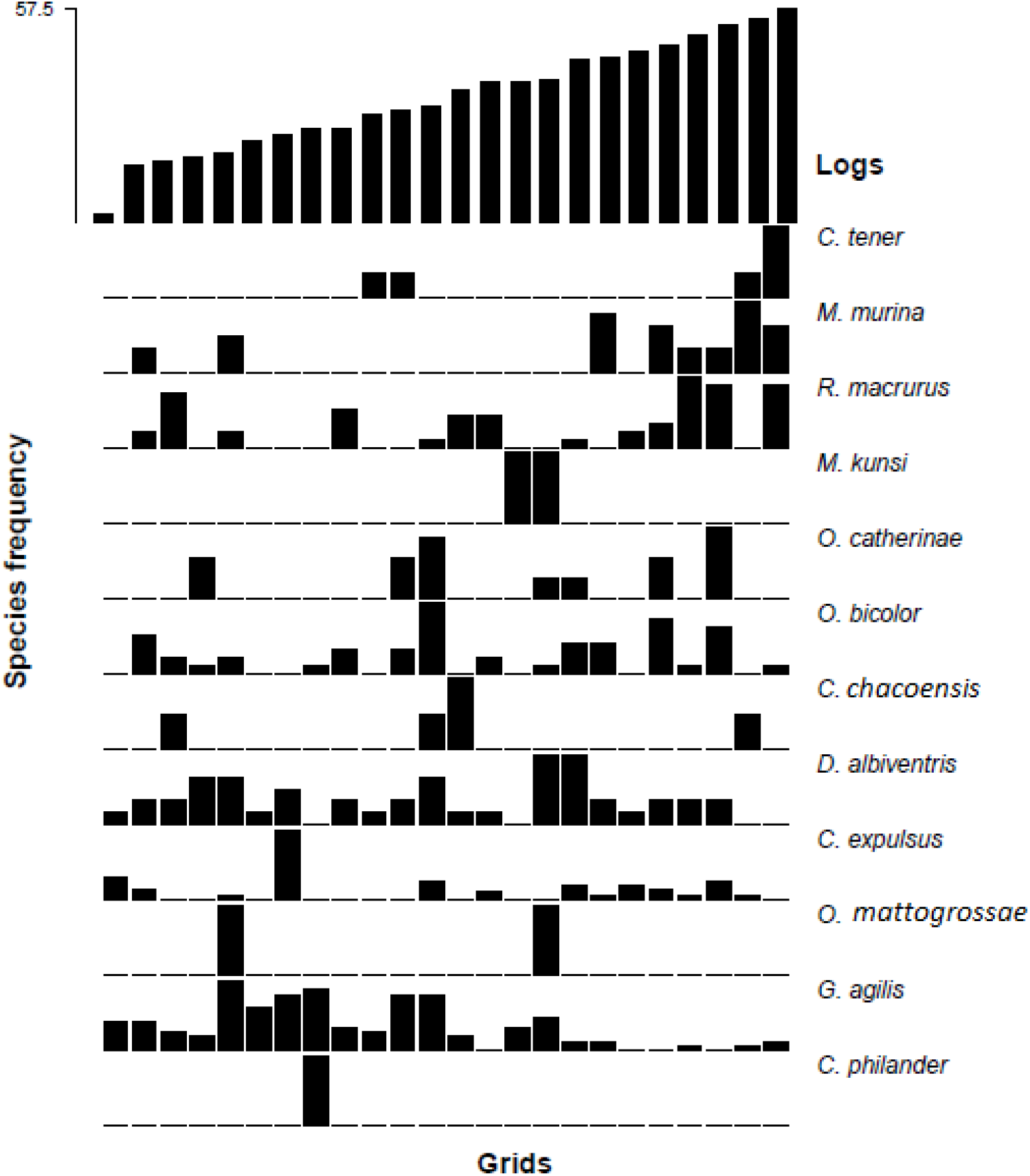
Direct ordination of small mammal species’ frequency associated with no. of fallen logs in fragmented landscape studied.

**S2 Fig.**
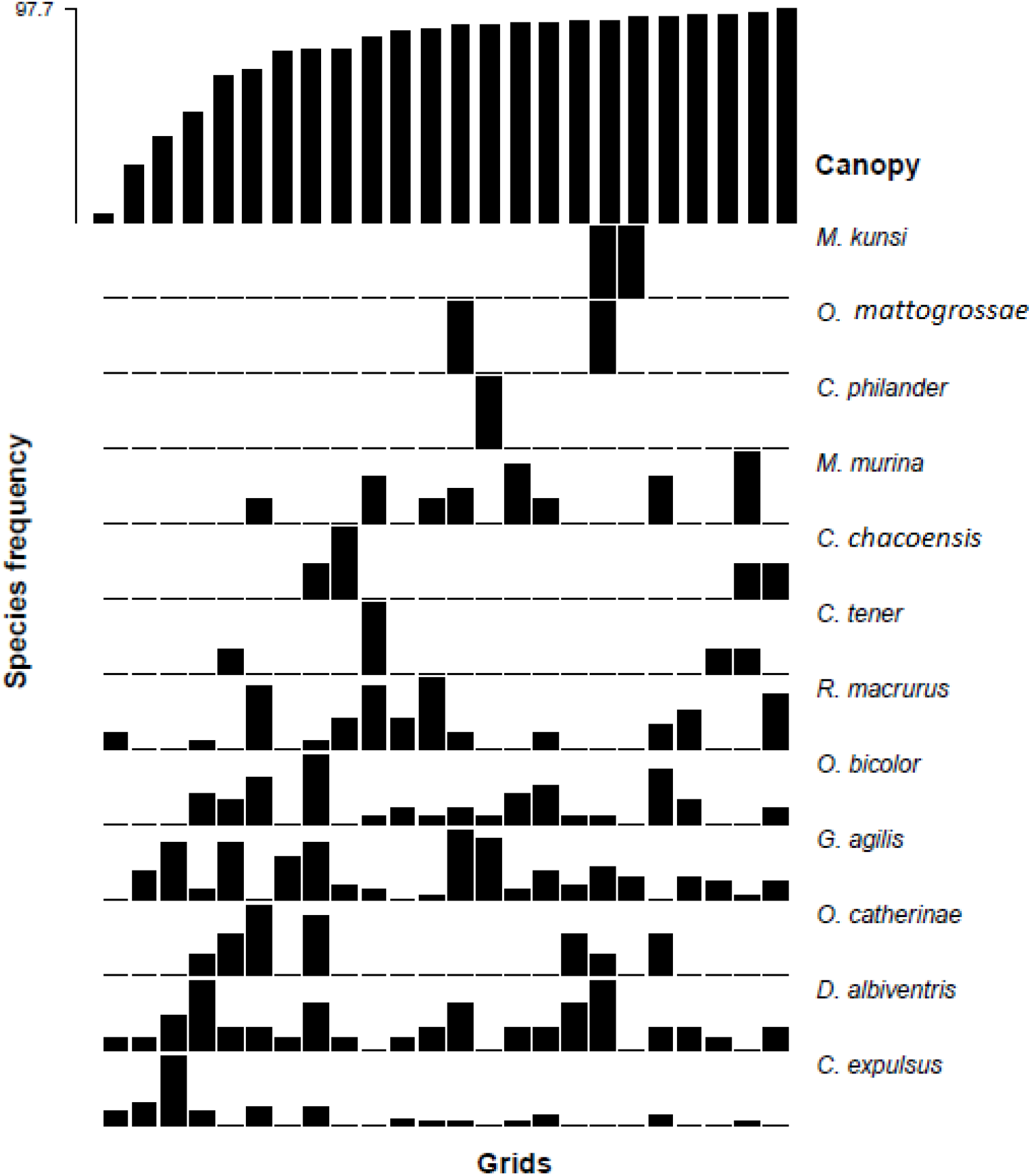
Direct ordination of small mammal species’ frequency associated with canopy cover in fragmented landscape studied.

**S3 Fig.**
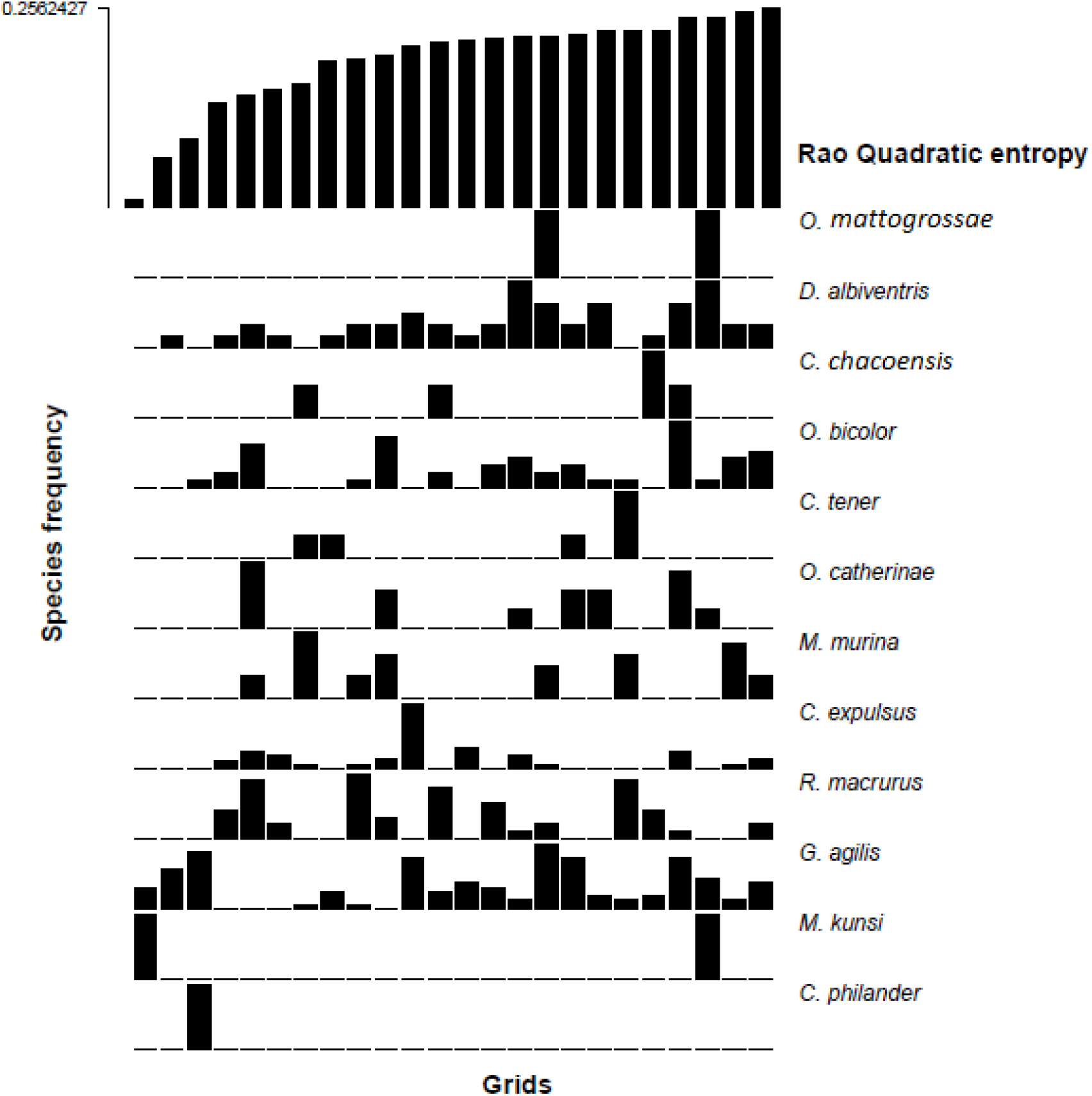
Direct ordination of small mammal species’ frequency associated with Rao Quadratic Entropy in fragmented landscape studied.

